# microRNA-Mediated Effects of Caloric Restriction on Trophoblast Invasion and Angiogenic Signaling in a Mouse Model of Fetal Growth Restriction

**DOI:** 10.1101/2025.02.05.636641

**Authors:** James R. Bardill, Caitlin R. Eason, Holly Wood, Courtney Breckenfelder, Lauren T. Gallagher, Madison Crew, Anis Karimpour-Fard, Carmen C. Sucharov, Theresa Powell, Clyde J. Wright, S Christopher Derderian

## Abstract

Trophoblast invasion is essential for normal placentation, with failure resulting in a fetal growth restriction (FGR) phenotype. Utilizing a calorie-restricted mouse model, we report progressive epigenetic, molecular, and phenotypic placental changes throughout gestation. Following maternal caloric restriction initiated at E9, we observed a significant reduction in fetal and placental weights beginning at E12.5, with persistent growth restriction at E14.5, E16.5, and E17.5. Immunohistochemistry of the decidual invasion site at E17.5 demonstrated reduced 1) decidual depth, 2) trophoblast invasion distance, and 3) trophoblast quantity within the decidua. Preceding these phenotypic changes, RT-qPCR revealed downregulation of trophoblast invasion and angiogenesis genes, including *MMP2, MMP9, EFNA1, Rac1, Rras, ASCL2, TRAP2C, Prl7b1, VEGFa, VEGFb, PDGF*, and *AKT3*, beginning as early as E14.5. Notably, microRNA sequencing at E12.5, prior to these transcriptional changes, identified significant upregulation of *miR-503-5p*, a predicted inhibitor of several of these pathways. The summation of these observations suggests *miR-503-5p* may be an early driver of placental dysfunction in FGR, linking maternal malnutrition to impaired trophoblast invasion and angiogenesis. These findings provide insight into the molecular mechanisms underlying placental insufficiency and highlight *miR-503-5p* as a potential therapeutic target for improving pregnancy outcomes in FGR.

## INTRODUCTION

Fetal growth restriction (FGR) is a pregnancy complication characterized by impaired fetal growth, with significant health consequences for the affected offspring.^1^ Successful placentation is necessary to develop a maternal-fetal interface that supplies adequate nutrients and oxygen to the developing fetus. This process requires trophoblast invasion into the decidua where they remodel the maternal spiral arteries and promote vasodilation. In FGR, placental histology often demonstrates small shortened, villi, with reduced trophoblast proliferation and invasion.^1–3^

A mouse caloric restriction model, like the one used in this report, is an invaluable tool for mapping the complex mechanisms underlying FGR. While trophoblast invasion is shallower in mice compared to humans, this model enables precise manipulation and analysis of placental development, which is not feasible in human pregnancies.^4^ Additionally, ∼60% of microRNA (miRNA) loci are conserved between mice and humans, a significantly higher percentage than in other species, making this model particularly relevant to study epigenetic regulation.^5,6^ Although FGR arises from diverse risk factors, maternal malnutrition remains a leading—and notably preventable—contributor worldwide. Thus, a caloric restriction model not only closely mirrors human pathology but also provides critical mechanistic insights, supports controlled experimentation, and holds strong translational potential.^7^

miRNAs are small non-coding RNAs that function as post-transcriptional regulators of gene expression. By binding to target messenger RNAs (mRNAs), miRNAs can inhibit translation or promote mRNA degradation, thereby regulating protein expression. Increasing evidence suggests that miRNAs have major roles in placental development, including trophoblast invasion and placental angiogenesis.^8–13^ Both up- and down-regulated miRNAs have been identified in placentas affected by FGR at the time of delivery; however, few studies have investigated their role in the pathogenesis of FGR.^14,15^

To address a key knowledge gap critical to *in utero* therapeutics, we utilize a calorie-restricted mouse model of FGR to investigate the dynamic interplay between miRNAs and mRNAs in regulating trophoblast invasion into the decidua and placenta angiogenesis across multiple gestational timepoints. By integrating miRNA with target gene expression profiles and histological findings, we aim to provide a comprehensive understanding of the molecular mechanisms impairing placentation in FGR.

## Materials and Methods

### Sex as a Biological Variable

Our study exclusively examined female mice because the disease modeled is only relevant in females.

### FGR Mouse Model

All animal experiments were conducted at the University of Colorado Anschutz medical campus with approved IACUC protocol 1181. Timed matings were performed, and vaginal plugs were screened for daily. Pregnant C57BL/6 mice were provided ad libitum access to food between E1–E8.5. From E8.5 until tissue procurement, dams received either a 50% caloric restricted diet (FGR) or continued ad libitum access (controls) as previously described.^16^ Mouse placentas were harvested from a minimum of three different dams within the caloric restricted and control groups, with a minimum of 5 placentas per pregnant dam analyzed per group at each timepoint. Placentas from each dam were randomly assigned for qPCR or histological analysis. Placentas for RNA analysis were snap-frozen in liquid nitrogen and stored at −[80 °C until further use. Placentas for histological analysis were fixed in 10% neutral buffered formalin overnight, followed by 70% ethanol at 4°C until processing for histology.

### RNA Extraction

Total RNA including miRNA was extracted from mouse placenta samples using a bead homogenizer and Trizol LS reagent (ThermoFisher, 10-296-028)/chloroform method. RNA concentration was measured using the NanoDrop 2000c spectrophotometer (Thermofisher Scientific, Waltham, MA, USA). If necessary, a RNA clean and concentrator kit (Zymo Research R1019, Irvine, CA, USA) was used to remove any contaminating DNA, phenol, and concentrate samples to improve 260/280 (1.9-2.0) and 260/230 ratios (2.0-2.2).

### Poly(A) RNA Sequencing

A set of E12.5 samples (5 control and 5 FGR) underwent Poly-A RNA sequencing by LC Sciences. The total RNA quality and quantity were analyzed with a Bioanalyzer 2100 and RNA 6000 Nano LabChip Kit (Agilent, CA, USA) with RIN number >7.0. Paired end 2 × 150bp sequencing on an Illumina Novaseq™ 6000 was performed at LC Sciences (Houston, Texas, USA) following the vendor’s recommended protocol. A cDNA library constructed from the pooled RNA from mouse placentas was sequenced with Illumina NovaseqTM 6000. Reads were aligned to the mouse reference genome and then mapped the reads to the reference genome. Genes differential expression analysis was performed by DESeq2 software between two different groups (and by edgeR between two samples).

### miRNA Sequencing

The same set of E12.5 mouse placentas RNA samples underwent miRNA sequencing. All extracted RNA was used in the library preparation following Illumina’s TruSeq-small-RNA-sample preparation protocols (Illumina, San Diego, CA, USA). Quality control analysis and quantification of the DNA library were performed using Agilent Technologies 2100 Bioanalyzer High Sensitivity DNA Chip. Single-end sequencing (1 µg of total RNA) was used to prepare small RNA libraries according to TruSeq Small RNA Sample Prep Kits (Illumina, San Diego, USA) protocol. Single-end sequencing 50 bp was performed on an Illumina Hiseq 2500 (Hangzhou, China). Unique sequences with length in 18∼26 nucleotide were mapped to *Mus musculus* precursors in miRBase 22.0 by BLAST search to identify known miRNAs and novel 3p- and 5p-derived miRNAs. Differential expression of miRNAs based on normalized deep-sequencing counts were analyzed by selectively using Fisher exact test, Chi-squared test, Student *t* test, or ANOVA based on the experiments design. The significance threshold was set to be 0.01 and 0.05 in each test.

### PANTHER and Ingenuity Pathway Analysis (IPA) of mRNA-miRNA Target Genes

PANTHER was used to analyze mRNA expression profiles to identify significantly altered biological pathways in the RNA sequencing dataset. mRNA and miRNA sequencing datasets were combined in a core analysis using IPA to identify significantly expressed canonical pathways in the datasets with predicted up and down regulation.

### RT-qPCR

Following RNA extraction, cDNA was synthesized, and relative mRNA and miRNA levels were evaluated by quantitative RT-qPCR. Quantification was performed using the cycle threshold (ΔΔCt) method. Primers for specific miRNAs as well as mRNAs of interest were used to compare relative levels of expression between caloric-restricted and control placentas. For mRNA, GAPDH was used as the normalizing gene. For miRNA, U6 was used as the normalizing gene.

### Histology/Immunohistochemistry

Placentas and their associated decidua were embedded in paraffin and cut at 5μm thickness. After deparaffinization and rehydration, antigen retrieval was performed with sodium citrate buffer (pH 6.0) using the IHC-TEK Epitope Retrieval Steamer. Sections were treated with a peroxidase blocking step and co-stained using IHC-Tek Antibody Diluent (Fisher NC9953741) at 1:300 overnight at 4°C. Secondary staining was performed at 1:500 in IHC-Tek Antibody Diluent. Images were obtained under a Leica DMi8 and analyzed using ImageJ. Primary antibodies used included Anti-Cytokeratin 8 (rabbit, EP1628Y, Abcam) and Mouse Endomucin (goat, AF4666, R&D Systems). Secondary antibodies were Anti-Goat 594 and Anti-Rabbit 488. Placenta zones (labyrinth, junctional zone, and decidua) were measured for total area and depth of each zone and CK8 positive cells were counted in the decidua.

### Statistical Analysis

Statistical analyses are performed using Prism GraphPad version 6 (GraphPad Software, Inc., La Jolla, CA). The alpha value p<[0.05 was considered statistically significant.

## RESULTS

### Caloric Restriction in a Pregnant Mouse Model Reduces Fetal and Placental Weights

To evaluate the efficacy of our caloric restriction mouse model in inducing FGR, we assessed placental and fetal growth trajectories across gestation. Placental and fetal weights were measured in both FGR and control groups at five embryonic time points: E10.5, E12.5, E14.5, E16.5, and E17.5. Placental weights in the FGR cohort were significantly lower as early as E12.5, just four days after the initiation of caloric restriction (**Figure 1A**). In contrast, fetal weights did not show a significant decrease until E16.5 (**Figure 1B**), suggesting that the placenta initially compensates to sustain fetal growth until its functional capacity is exceeded. **Figure 1C** provides representative pictures of placental and fetal weights throughout gestation. These anatomic observations validate the efficacy of our caloric restriction model to induce FGR, as evidenced by the consistent and significant impairment of placental and fetal growth.

**Figure 1.**
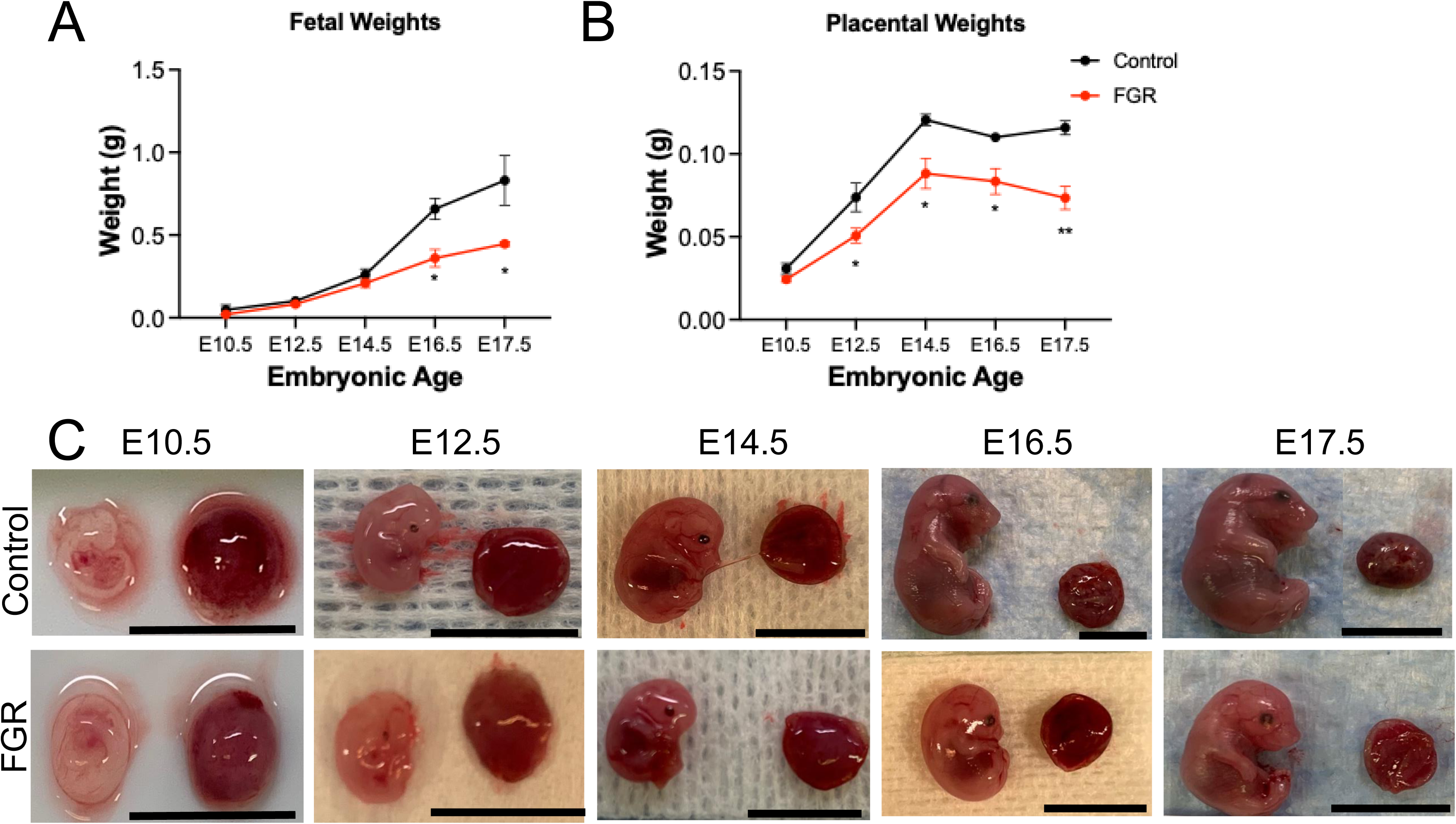
Caloric *restriction mouse model results in reduces placental and fetal weights.* *(A)* Placenta weights are significantly reduced in fetal growth restricted (FGR) mice beginning at E12.5. (B) *Fetal weights* are significantly reduced in FGR mice beginning at *E16.5*. (C) Representative images of control and FGR mouse placentas and fetuses throughout gestation illustrating reduced sizes within the FGR cohort (C). * = p<0.05, ** p<0.01 determined by student’s t-test. Scale bar in each image represents 1 cm.

### Trophoblast Invasion Assessed by Immunohistochemistry

Trophoblast invasion was assessed by IHC of placentas and their adjacent decidua throughout gestation. Invasion sites procured from pregnancies associated with caloric restriction had a reduced decidual thickness and fewer trophoblasts within the decidual zones when compared to controls, an observation evident as early as E14.5 (**Figure 2A**). We quantified the depth of the decidua, the number of CK8+ trophoblasts within the decidua, and the depth of CK8+ trophoblast invasion into the decidua. Decidual depth was significantly reduced and fewer trophoblasts were observed within the decidua in our FGR cohort as early as E14.5 (**Figure 2B-2C**). Lastly, there was a significant reduction in the depth of trophoblast invasion through the decidua in FGR placentas at E16.5 and E17.5 (**Figure 2D**). No significant differences in the endothelial marker, endomucin, were observed within the three placental zones between FGR and control placentas.

**Figure 2.**
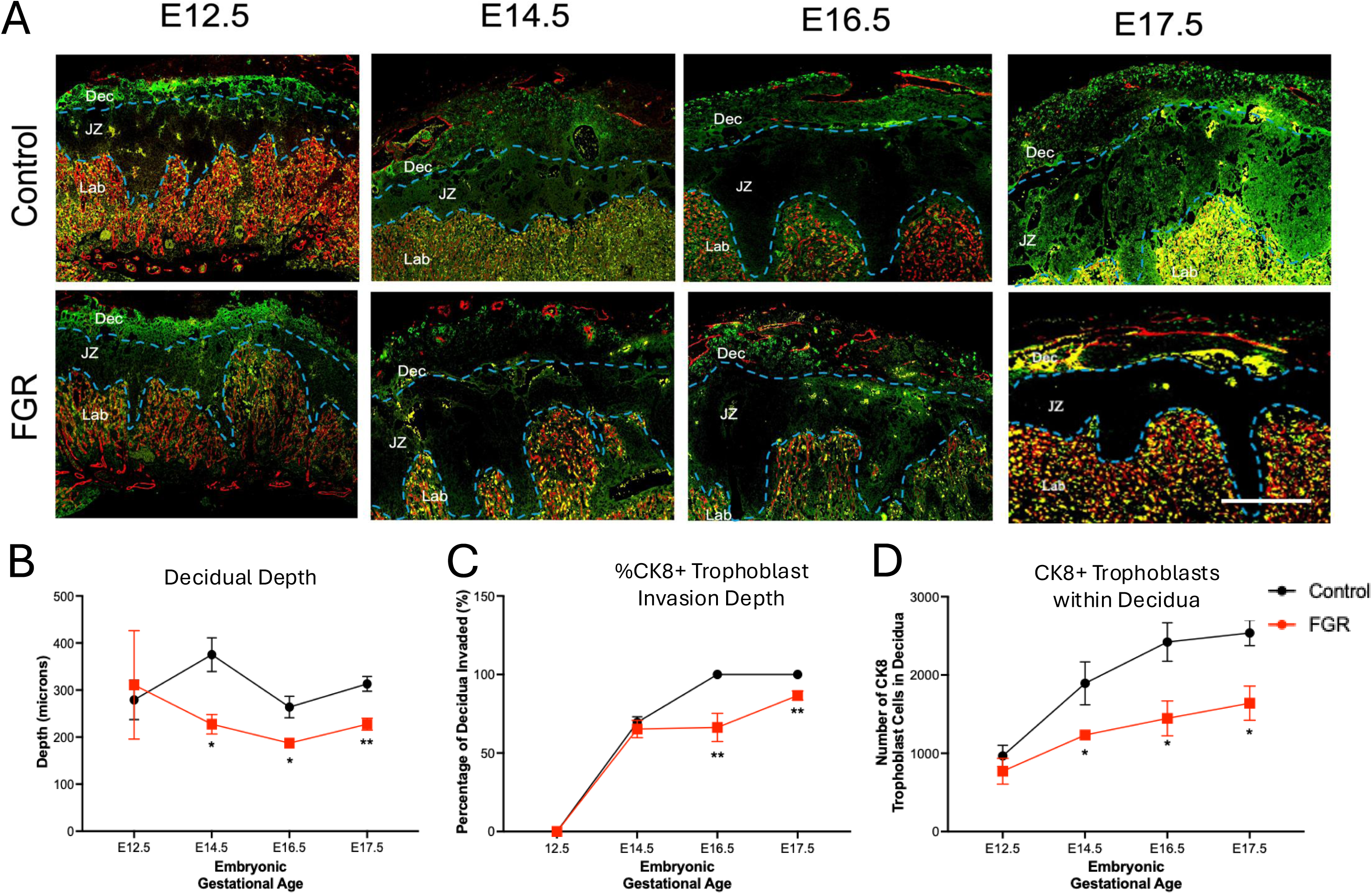
Caloric restriction results in impaired decidual trophoblast invasion. *(A)* Immunohistochemistry of invasion sites obtained using Thunder Imager with a 20X magnification. Endothelial cells are stained with Endomucin (red), and trophoblasts are stained with Cytokeratin 8 (green). (B) placenta decidua depth across gestation showing a significant reduction in FGR decidua compared to control decidua, (C) percentage of CK8+ cells that reach full invasion depth of the decidua, and (D) total number of CK8 cells in the decidua. Lab: labyrinth, JZ: junctional zone, Dec: decidua. Scale bar represents 500 mm. *p<0.05 and **p<0.01 by students t-test.

### Downregulation of Genes Associated with Trophoblast Invasion and Placental Angiogenesis

To assess the impact of caloric restriction on trophoblast invasion, we examined the expression of key genes involved in this process using RT-qPCR. mRNA expression levels of *EFNA1, MMP2*, *MMP9*, *ASCL2*, *Rac1*, *Rras*, *TRAP2C,* and *Prl7b1* were measured in placentas from FGR and control cohorts at gestational timepoints: E10.5, E12.5, E14.5, E16.5, and E17.5 (**Figure 3**). A majority of genes we analyzed were downregulated in our FGR model as early as E16.5. Only *EFNA1* was significantly downregulated at an earlier timepoint (E14.5). Because trophoblast invasion and placental angiogenesis are complementary processes altered in FGR, we also analyzed gene expression levels of key proteins associated with cell invasion and endothelial angiogenesis, namely *VEGFa, VEGFb, PDGF* and *AKT3.* Our RT-qPCR analysis indicated that *VEGFa* and *AKT3* were downregulated in FGR placentas as early as E14.5 with the downregulation of *VEGFb* and *PDGF* lagging by two days (**Figure 4**).

**Figure 3.**
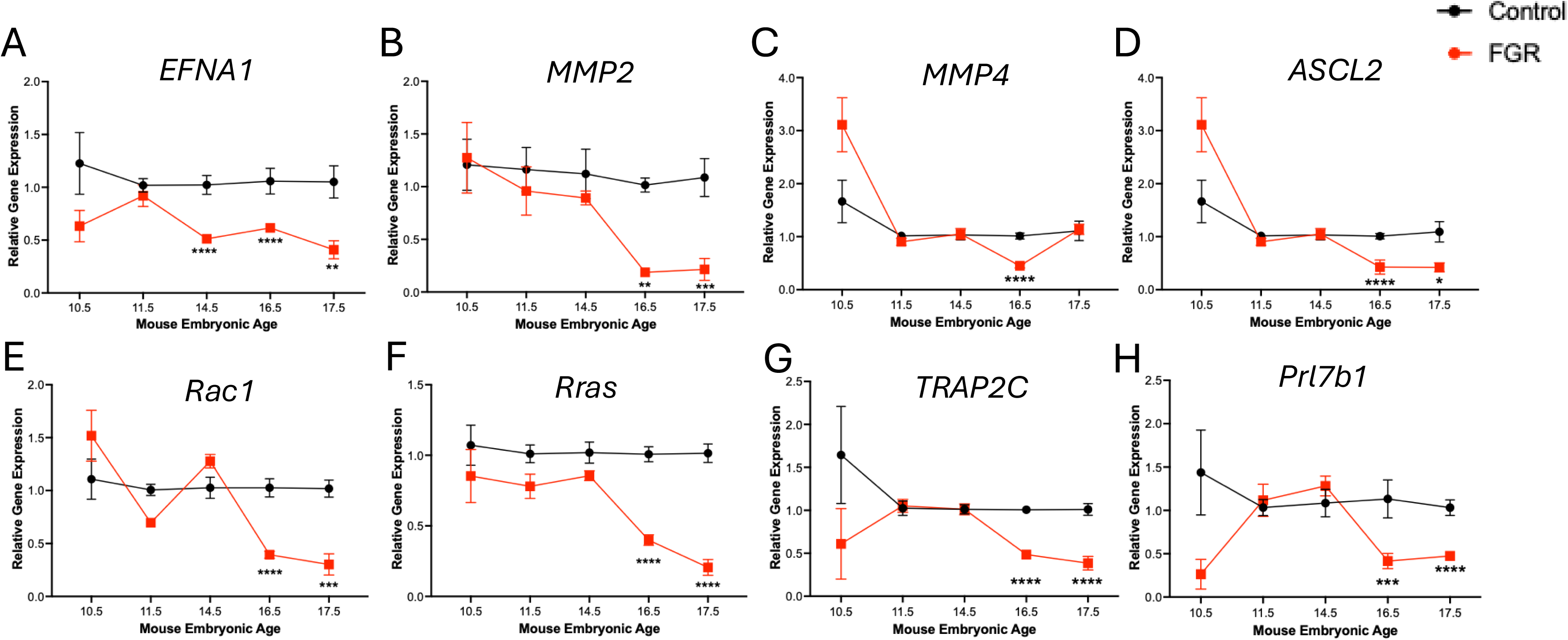
RT-qPCR comparing fetal growth restriction (FGR) and control mouse placenta trophoblast invasion genes throughout gestation. (A) *ephrin-A1 (EFNA1)*; (B) *matrix metalloproteinase-2 (MMP2)*; (C) *matrix metalloproteinase-9 (MMP9*; (D) *Achaete-scute-like 2 (ASCL2)*; (E) *Ras-related C3 botulinum toxin substrate 1 (Rac1)*; (F) *ras-related protein (Rras)*; (G) *transcription factor AP-2 gamma (TRAP2C)*; and (H) *prolactin family 7, subfamily b, member 1 (Prl7b1)*. *p<0.05; **p<0.01; ***p<0.001; ****p<0.0001 by students t-test.

**Figure 4.**
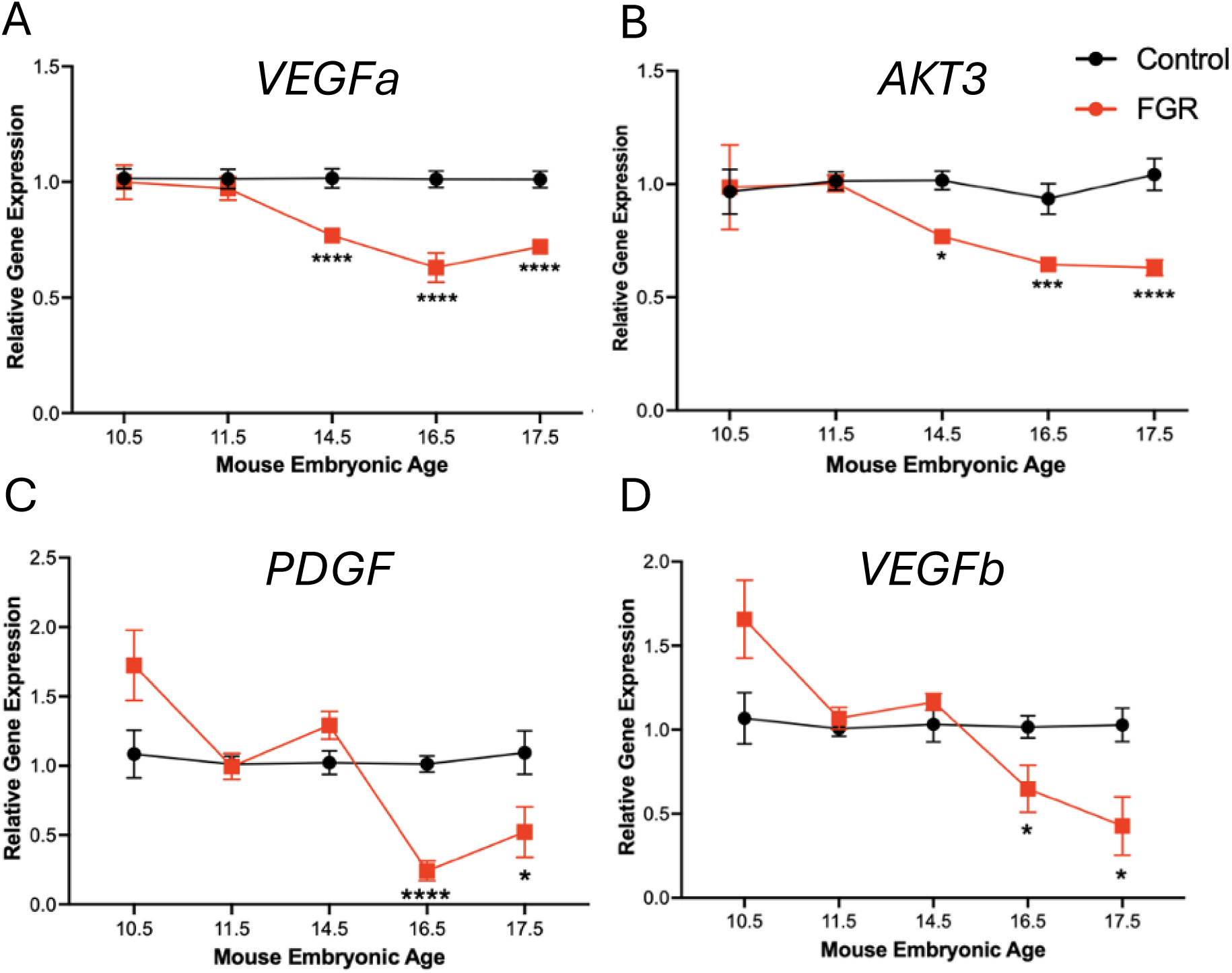
RT-qPCR comparing fetal growth restriction (FGR) and control mouse placenta angiogenic genes throughout gestation. (A) *VEGFa,* (B) *AKT3,* (C) *PDGF* and (D) *VEGFb*. *p<0.05; **p<0.01; ***p<0.001; ****p<0.0001 by students t-test.

### Early microRNA Changes Resulting from Maternal Caloric Restriction

To determine the early effects of caloric restriction on miRNAs, we sequenced whole placentas at E12.5. A total of 1,336 miRNAs were identified within this analysis. Among them, 63 were differentially expressed within the FGR cohort compared to control cohort (31 upregulated and 32 downregulated). A volcano plot and heatmap of significantly expressed miRNAs that target mRNAs of proteins associated with invasion and angiogenesis are presented in **Figure 5**. The top 20 significantly expressed miRNAs sorted by highest counts are displayed in the **Table**. PANTHER analysis of RNA-sequencing from E12.5 placentas revealed significant downregulation in angiogenesis pathways, including *VEGF, EGF, PDGF,* and endothelin signaling (**Figure 6**).

**Figure 5.**
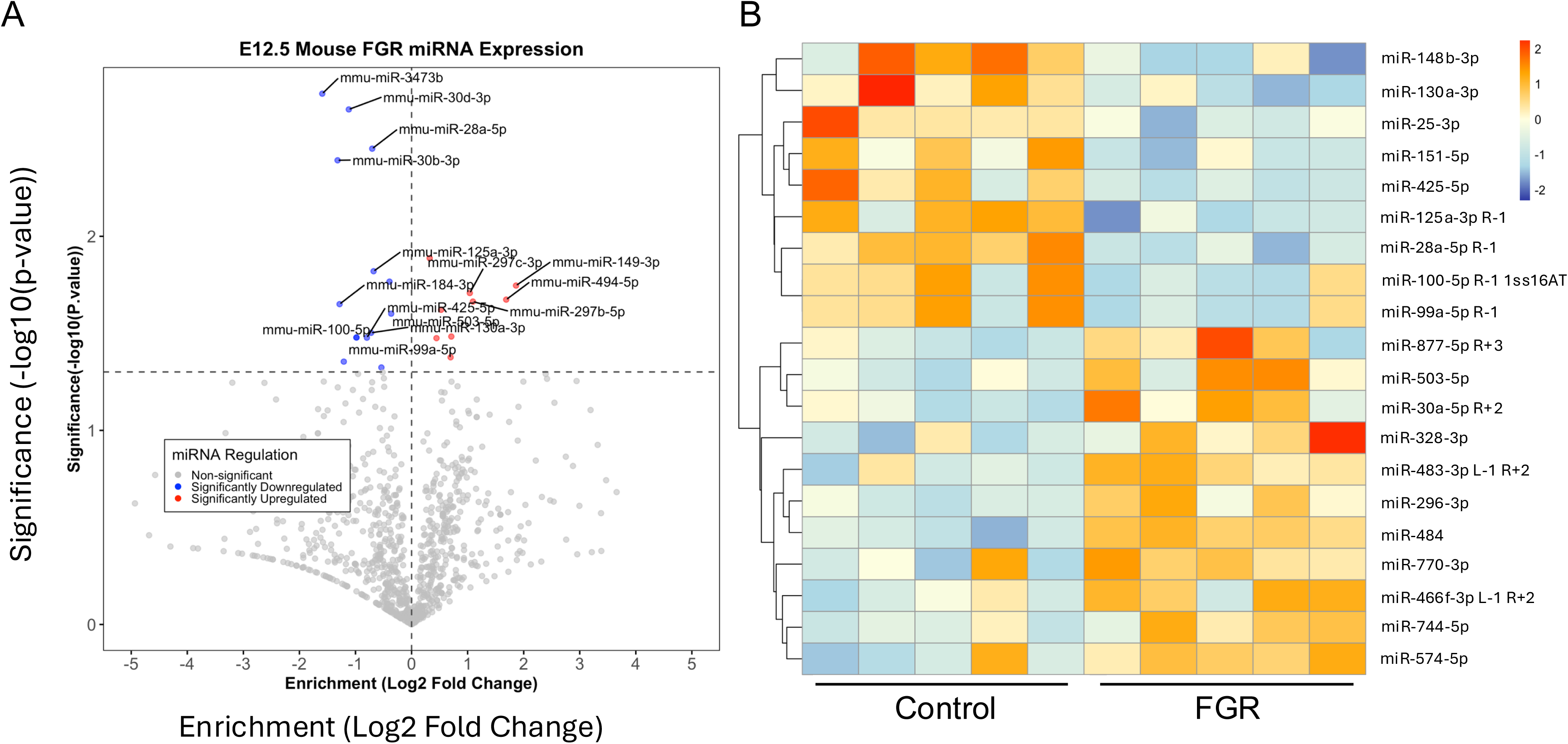
microRNAs with significantly different levels of expression between fetal growth restricted (FGR) and control placentas. (A) Volcano plot of microRNA sequencing results from whole placental tissue harvested at E12.5. Blue dots denote significantly downregulated microRNAs and red dots denote significantly upregulated microRNAs. (B) Heatmap of differentially expressed microRNAs that are known to target mRNAs encoding proteins involved in invasion or angiogenesis, comparing FGR and control placentas.

**Figure 6.**
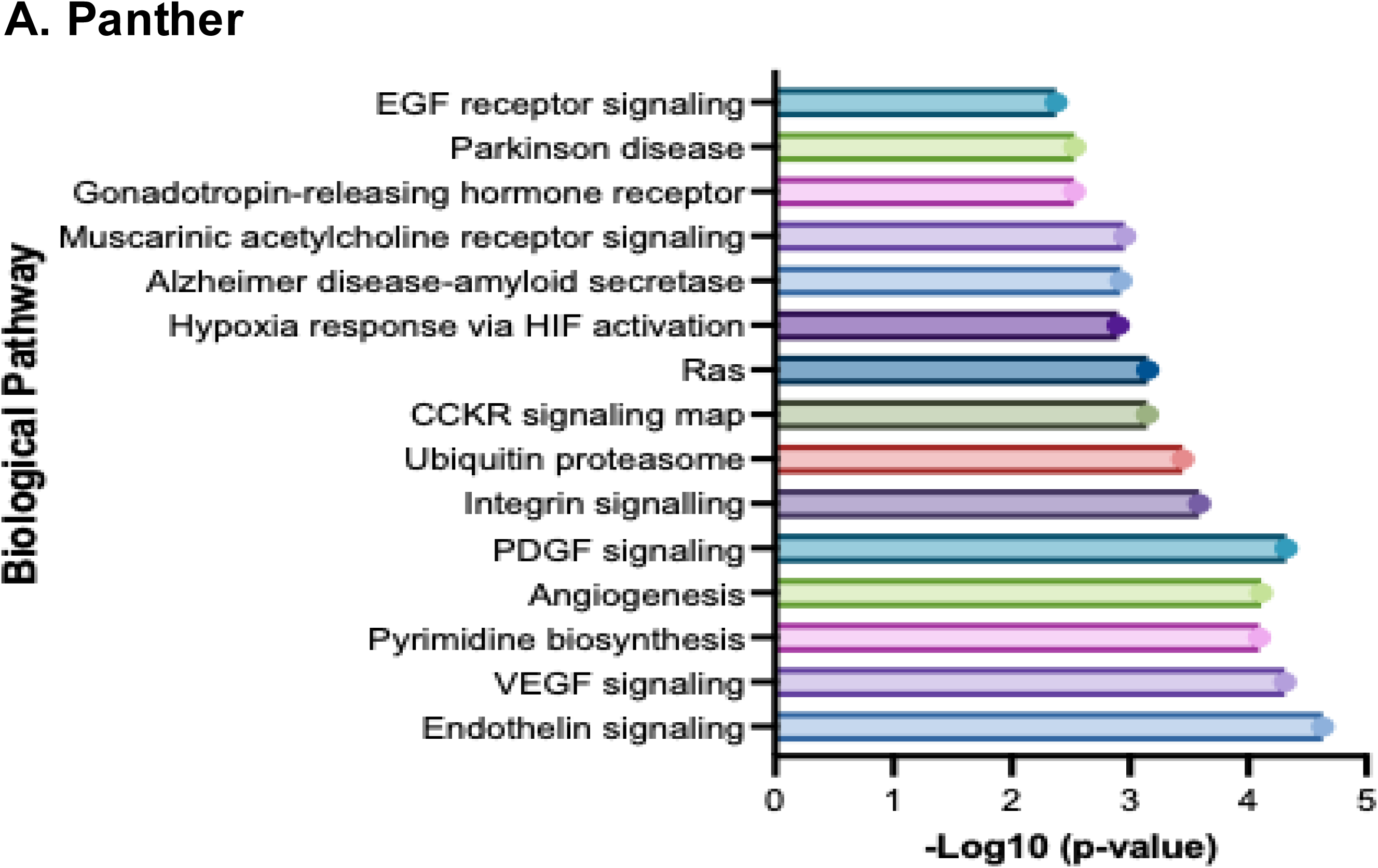
microRNA sequencing analysis. (A) PANTHER analysis of E12.5 mouse placental mRNA sequencing identifying multiple significant altered biological pathways in fetal growth restricted placentas. (B) KEGG pathway analysis of microRNA sequencing.

**Table.**
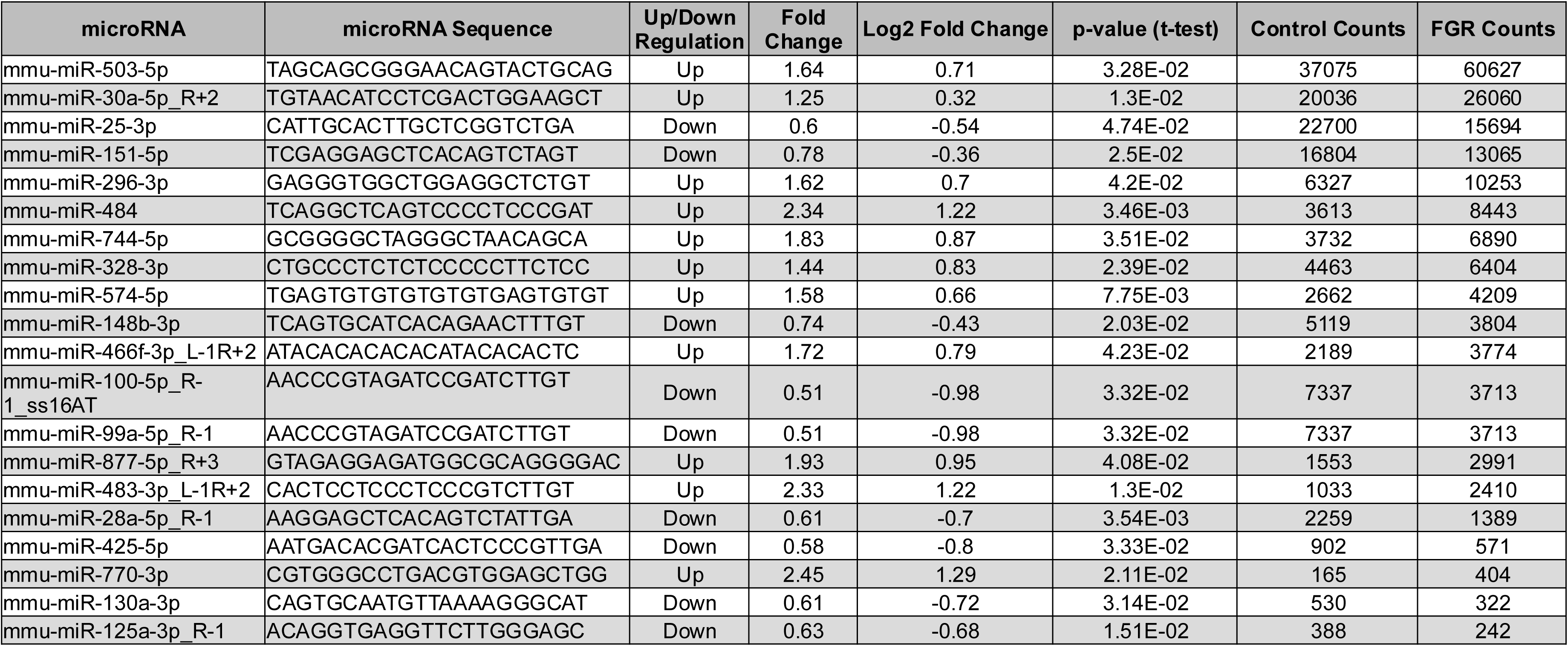
Top 20 significantly expressed miRNAs. Fetal growth restricted (FGR) and control placentas analyzed at E12.5. microRNAs listed in order from highest to lowest counts. Count values represent the mean value of five samples per group.

IPA of the combined E12.5 miRNA and mRNA sequencing data revealed significant alterations in several key pathways in FGR placentas compared to controls. Notably, we observed a downregulation of metabolic pathways, including oxidative phosphorylation and electron transport chain, coupled with an upregulation of mitochondrial dysfunction pathways (**Figure 7**). These findings reflect the impact of caloric restriction on placental energy metabolism. Additionally, pathways related to RNA processing and translation were also predicted to be downregulated, potentially reflecting the regulatory influence of miRNAs on gene expression. Conversely, pathways related to mitochondrial dysfunction and degradation of the extracellular matrix were predicted to be upregulated (**Figure 7**).

**Figure 7.**
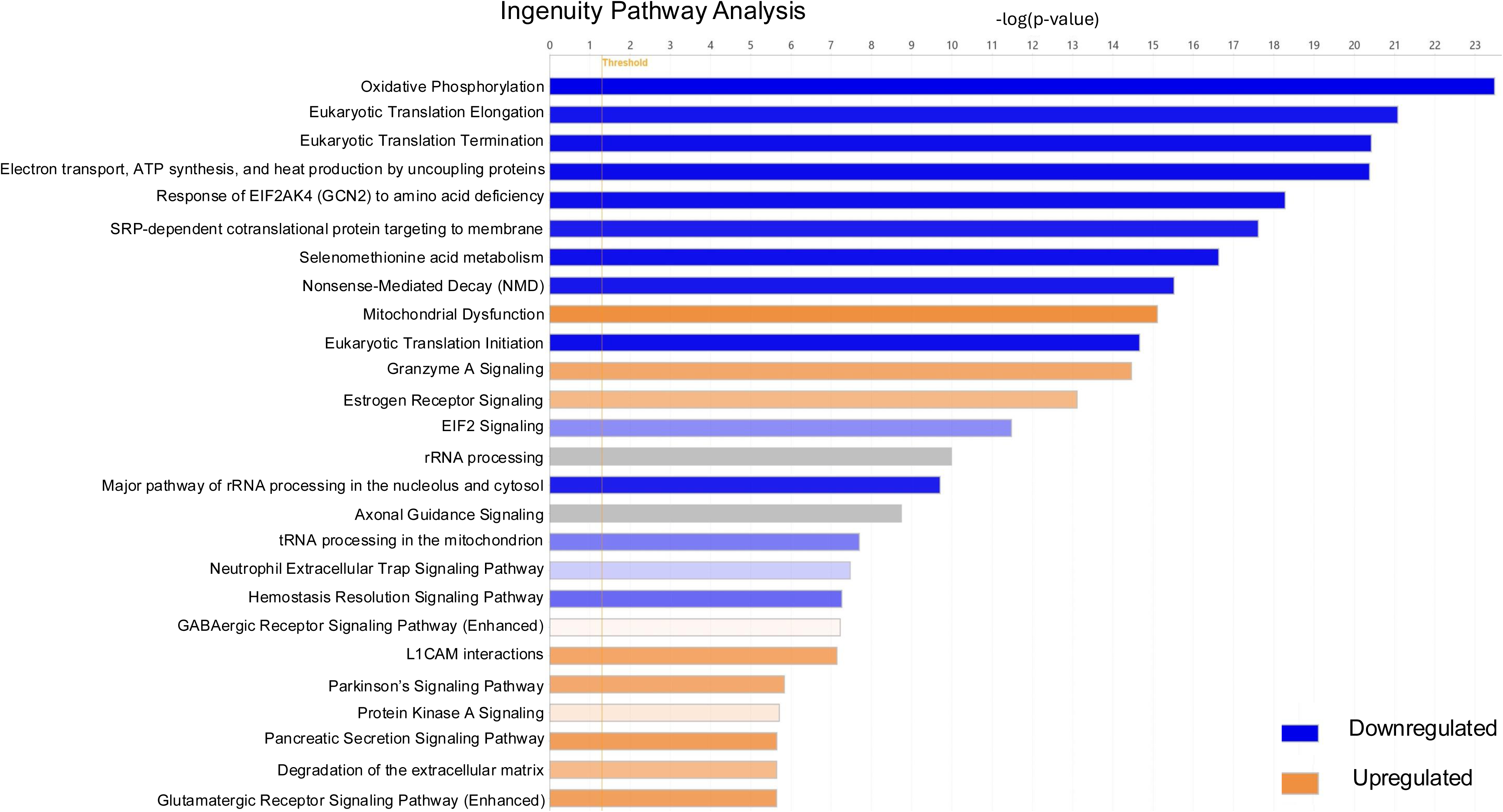
Ingenuity Pathway Analysis (IPA) of E12.5 whole placental mRNA and microRNA. Blue rows represent downregulated pathways while orange rows represent upregulated pathways.

Among the highly expressed miRNAs upregulated in our E12.5 caloric-restricted placentas from miRNA sequencing analysis, miR-503-5p was notable for its predicted targeting of genes involved in invasion and angiogenesis. In fact, IPA predicted miR-503-5p to downregulate genes involved with angiogenesis, proliferation, and cell migration, including *VEGFa, WNT4*, and *AKT3* (**Figure 8A**). miRT-qPCR verified the significant upregulation of miR-503-5p in E12.5 placenta samples seen in the miRNA sequencing (p<0.01, **Figure 8B**). From our longitudinal RT-qPCR analysis, *VEGFa* and *AKT3* were found to be downregulated as early as E14.5. However, *WNT4* gene expression was not significantly different between FGR and control samples (data not shown). Thus, two of the three predicted targets of miR-503-3p (*VEGFa and AKT3*) were downregulated two days after miR-503-5p was shown to be upregulated. These targets likely serve as key mediators influencing additional canonical pathways essential for the coordinated angiogenic and invasive processes required for placentation.

**Figure 8.**
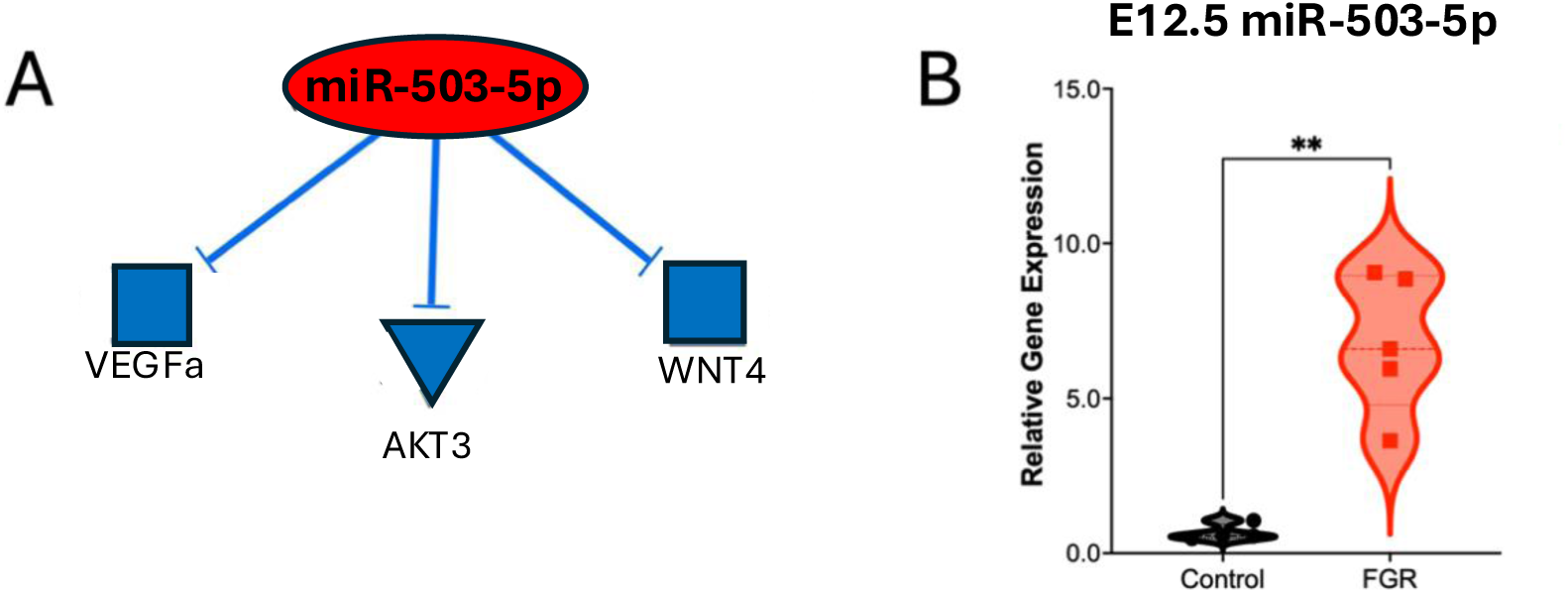
miR-503 validation and gene expression. (A) Predicted targets of miR-503-5p by Ingenuity Pathway Analysis (IPA). Red denotes increased expression and blue denotes predicted downregulation. (B) miRNA RT-qPCR verified miRNA sequencing results of miR-503-5p with significant upregulation in FGR placentas as E12.5.5. n = 5 per cohort from three separate pregnancies.

## DISCUSSION

This analysis demonstrates the significant impact of maternal nutrition on placental development and trophoblast invasion. We observed substantial reductions in both fetal and placental weights, with placental growth impairment occurring earlier than FGR, suggesting an initial compensatory phase that is ultimately unsustainable. Transcriptomic analysis revealed widespread alterations in gene expression, including the downregulation of key pathways involved in trophoblast invasion and placental angiogenesis. Notably, we identified *miR-503-5p* as a highly expressed and upregulated miRNA in FGR placentas, with predicted targets *VEGFa* and *AKT3* showing subsequent downregulation.^17^ These findings suggest a mechanistic link between anti-invasive miRNA upregulation and impaired placentation. Histological analysis further confirmed a reduction in trophoblast invasion and decidual thickness, reinforcing the hypothesis that caloric restriction disrupts essential placental processes. Collectively, our results provide novel insights into the molecular mechanisms driving FGR, highlighting potential opportunities to study therapeutics.

### Caloric Restriction in Pregnant Mice as a Model of FGR

Although FGR has diverse risk factors, malnutrition accounts for a significant proportion of cases. Globally, FGR affects an estimated 7%–15% of pregnancies, but its prevalence can rise to as high as 30% in low- and middle-income countries, emphasizing the importance of nutrition in fetal development.^18,19^ Multiple preclinical models to study FGR have been proposed. These include the ovine hyperthermia models, rodent uterine artery ligation models, and maternal nutrient-restriction models.^20–24^ We chose a maternal caloric restriction animal model because it has been shown to impair placental angiogenesis, disrupt trophoblast invasion, and alter fetal organ development, mirroring findings in human FGR.^25^ Nutrient-restricted models have also been shown to induce placental insufficiency, leading to altered umbilical blood flow dynamics and asymmetric FGR.^26^ This model also replicate key molecular mechanisms implicated in FGR, such as increased oxidative stress, dysregulated inflammatory responses, and epigenetic modifications affecting fetal development.^27^ Thus, nutrient-restriction-based models serve as a valuable tool to study the multifactorial pathogenesis of FGR and identifying potential therapeutic targets.

Our findings demonstrate that caloric restriction leads to reduced fetal and placental weights, validating the FGR phenotype. We were intrigued by the observation that fetal weight loss lagged that of placenta. It is probable that although the placenta does not have the nutrients necessary to its own proliferate, it selflessly prioritizes nutrient delivery to the fetus. The finding that reduced placental weights may precede the diagnosis of FGR has been reported in several human studies.^28–30^ Although this was not the primary focus of our study, we found this observation compelling and consider it a significant finding. The suggested compensatory response warrants further investigation.

Our mRNA-miRNA sequencing analyses at E12.5 not only provides further validation of this model, but through advanced bioinformatics tools such as PANTHER, KEGG, and IPA, we identified a reduction in key metabolic pathways, including oxidative phosphorylation and electron transport, coupled with an increase in mitochondrial dysfunction. This metabolic stress imposed by nutrient restriction aligns with previous findings in mice at E17.5 demonstrating alterations in inflammatory and hypoxia-related pathways.^31^ We speculated that caloric restriction induces early and sustained stress at the cellular level which precedes downregulation of molecular mechanisms critical for placentation, specifically trophoblast invasion and placental endothelial angiogenesis.

### Mouse Model of FGR Impairs Key Genes in Trophoblast Invasion

Following the initial insult resulting from maternal caloric restriction (oxidative stress coupled with altered levels of miRNA expression), genes critical to trophoblast invasion were downregulated, with the earliest effects occurring at E14.5, five days after caloric restriction. These alterations included reduced expression of *ASCL2*, a transcription factor critical for trophoblast differentiation and function, suggests a broad impact of caloric restriction on trophoblast development.^32,33^ In addition, the downregulation of *EFNA1*, *Rac1*, and *Rras* genes involved in cell signaling and cytoskeletal rearrangement, support the notion of impaired trophoblast migration and invasion in FGR.^34–36^

Other than molecules critical for cell motility, we found a significant reduction in MMP2 and MMP9 expression in FGR mouse placentas at E16.5 and E17.5. MMPs play crucial roles in cellular invasion. MMP2 and MMP9 are key enzymes involved in extracellular matrix degradation, a process essential for trophoblast invasion and proper placental development. This degradation facilitates trophoblast cell motility and invasion into the uterine tissue, enabling the establishment of a functional maternal-fetal interface. Downregulation of MMP2 and MMP9 activity can contribute to impaired trophoblast invasion, a hallmark of FGR. Studies in both *WNK1* knockout mice and *LAMC2*-overexpressing models demonstrate that altered *MMP2* and *MMP9* expression directly affects trophoblast invasion, highlighting the critical role of this enzyme in early placental development.^37^ I*n vitro* experiments suggest supplementation of MMP9 may even improve pregnancy outcomes.^38–40^ Conversely, knockdown of *MMP2* attenuates trophoblast invasion, underscoring the importance of this pathway in regulating placentation.^41^

Eph receptor-ephrin signaling pathways play a fundamental role in shaping cell behavior and tissue organization during embryogenesis. EFNA1 is a signaling molecule involved in cell-to-cell communication, guiding trophoblast cell migration and adhesion. EFNA1 directs trophoblast cells to the correct uterine location and promotes interactions with uterine epithelium. Our results demonstrated a decreased expression of *EFNA1* as early as E14.5. *EFNA1* is highly expressed in trophoblasts and promotes invasion of trophoblasts by engaging with the EphA2 receptor found on decidual epithelial cells.^36,42,43^

Eph-Ephrin signaling also activates Rac1, a vital molecular switch that orchestrates cell motility, adhesion, and cytoskeletal dynamics and is essential for trophoblast invasion. Rac1 further influences the expression and activity of MMPs, highlighting the complex interplay of signaling pathways regulating placentation. In our analysis, *Rac1* was significantly downregulated in FGR placentas beginning at E16.5. Preeclampsia, another pregnancy complication found to have reduced trophoblast invasion has also been shown to have significantly lower levels of *Rac1* and *MMP9* within placental samples.^34,44,45^ When *Rac1* is downregulated in cultured HTR-8/SVneo trophoblasts with a siRNA knockdown, trophoblast migration in reduced.^44,46^ Rras, a molecular switch crucial for cell movement and adhesion, was also downregulated in caloric-restricted placentas. While previous research has linked Rras to pathological cell behavior, this study is among the first to implicate it in placental dysfunction, specifically in the context of FGR.^47^

*ASCL2*, a transcription factor essential for trophoblast differentiation and the regulation of invasion-related genes, including MMPs, was significantly reduced in FGR placentas beginning at E16.5. This finding is consistent with previous studies demonstrating that altered *ASCL2* expression disrupts trophoblast differentiation and invasion, contributing to placental insufficiency and FGR.^48,49^ For example, a study in mice with a colony-stimulating factor mutation showed impaired *ASCL2* expression and trophoblast differentiation, while another study in rats linked ASCL2 loss to FGR and placental abnormalities.^50,51^ *TRAP2C*, a key transcription factor for trophoblast invasion and implicated in FGR,^52^ was also significantly downregulated in FGR placentas beginning at E16.5. Lastly, *Prl7b1*, which plays a role in placental angiogenesis and is highly expressed in invasive trophoblasts, was also reduced in our caloric restriction cohort beginning at this timepoint.^53,54^ Notably, alterations in other members of the prolactin family were observed in our E12.5 mRNA sequencing data, further supporting the role of disrupted prolactin signaling in FGR pathogenesis.

### Mouse Model of FGR Impairs Trophoblast Invasion

Our findings demonstrate that trophoblast invasion is clearly impaired in this FGR model, as evidenced by both the downregulation of key invasion-related genes and reduced trophoblast infiltration observed via IHC. This highlights the profound impact of caloric restriction on placental development, triggering a cascade of deleterious effects. Our results are consistent with previous studies showing that VE-cadherin, a protein integral to invasion, knockout mice exhibit shallow trophoblast invasion and reduced invasion depth and area.^55^ Similarly, our model revealed a significant reduction in decidual size and impaired trophoblast invasion beginning at E14.5. The observed decrease in decidual depth was unexpected, underscoring not only the fetal reliance on adequate nutrient supply but also the maternal requirement for sufficient resources to establish the microenvironment necessary to sustain pregnancy.

### Mouse Model of FGR with Potential miRNA Effects on Angiogenesis

Trophoblast growth, invasion, and uterine spiral artery remodeling are essential for proper placental development and fetal growth.^13^ Expanding upon previous studies that identified biological pathways and miRNAs involved in placental angiogenesis in FGR,^15,31^ this study incorporated an earlier gestational timepoint (E12.5) to capture the dynamic changes in miRNA and mRNA expression throughout placental development. In this study, we identified significant alterations in pathways critical for trophoblast invasion and spiral artery remodeling, including angiogenesis, PDGF, and integrin signaling. However, these processes extend beyond miRNA regulation alone—correcting FGR is not as simple as altering miRNA expression levels. For instance, natural killer cells contribute to spiral artery remodeling, angiogenesis, and trophoblast migration through cytokines such as IP-10 and VEGF, highlighting the multifaceted nature of these interactions.^56^ While our findings provide novel insights into factors contributing to FGR, they represent only part of the larger biological complexity.

A particularly compelling result was the significant upregulation of *miR-503-5p* in FGR placentas, accompanied by a corresponding reduction in key angiogenesis and invasion markers, including *VEGFa*, *VEGFb*, AKT3, and *PDGF. miR-503-5p* has been well-documented for its inhibitory effects on *VEGF* and has also been detected in extracellular vesicles of mouse trophoblasts, suggesting a potential role in cell-to-cell communication.^57–62^ Given that trophoblast invasion is intricately linked to angiogenesis in placental development and endometrial remodeling, the upregulation of *miR-503-5p* in FGR placentas points to a potential mechanism underlying impaired invasion and vascularization.^13^ Other miRNAs have also been implicated in FGR and preeclampsia; for example, increased expression of *miR-29b* has been shown to induce apoptosis, reduce trophoblast invasion and angiogenesis, and downregulate key targets such as *MCL1, MMP2, VEGFa,* and *ITGB1*.^62^ These findings reinforce the idea that miRNA-mediated regulatory networks play a crucial role in placental pathology and warrant further investigation as potential therapeutic targets.

### Conclusion

Our findings, demonstrating progressive impairment of trophoblast invasion throughout gestation in a caloric-restricted mouse model, underscore the detrimental impact of inadequate nutrient availability on placental development and function. This model recapitulates key features of FGR, including reduced fetal growth, placental insufficiency, and downregulation of molecular pathways critical for trophoblast invasion and placental angiogenesis. The identification of key genes, pathways, and miRNA/mRNA interactions altered at early gestational timepoints provides a valuable framework for understanding the pathogenesis of FGR. This work holds promise for developing novel therapeutic strategies to improve placental function, promote trophoblast invasion, and ultimately enhance fetal growth in FGR pregnancies.

## Author contributions

The manuscript was written with the contributions of all authors. All authors have approved the final version of the manuscript.

Conception or design of the work: JB, CD, CW

Acquisition, analysis, or interpretation of data: JB, CD, CE, HW, CB, MC, LG, CW, TH, AK, CS

Drafted the work or substantively revised it: JB, CD, CE, TH, CW, CS

All authors reviewed the manuscript.

## Acknowledgments

The authors thank the University of Colorado School of Medicine Morphology and Phenotyping Core for their assistance with tissue processing and immunohistochemistry. We would also like to acknowledge the University of Colorado Department of Surgery and Children’s Hospital Colorado for financial support to execute these studies.

## Disclosure statement

No potential conflict of interest was reported by the author(s).

## Funding

This work was supported in part by the Children’s Hospital of Colorado Ponzio Award

## Notes

**Conflict of Interest Statement** - The authors have declared that no conflict of interest exists.

### Competing Interest Statement

The authors have declared no competing interest.

